# Crop raiding as an emerging threat to giraffes: drivers and perceived effectiveness of countermeasures

**DOI:** 10.64898/2026.05.01.722334

**Authors:** Raymond O. Owino, Jessie Golding, Edwin L. Sangale, Abdullahi H. Ali, Jesse M. Alston

## Abstract

Giraffes, unlike other large mammals, typically pose minimal risk to humans, their animals, and crops, so they are traditionally not involved in human-wildlife conflict. Tree crops, however, are expanding across Africa, resulting in crop raiding by giraffes and retaliatory snaring, poaching, and killing of giraffes in response. The dynamics of giraffe crop raiding, however, are poorly understood, making effective interventions difficult to implement. To better understand key factors for humans and giraffes that mediate crop raiding, we used a multi-method approach to estimate giraffe abundance and activity, understand farmers’ perceptions and decisions, and test countermeasures around Garissa Giraffe Sanctuary in eastern Kenya. We hypothesized that 1) giraffe farm invasion would occur in dry seasons, 2) farms growing mangoes would be more likely to be invaded, 3) reducing invasion with only physical barriers would be less effective than adding behavior-based countermeasures, 4) perceptions would match giraffe activity and 5) countermeasure adoption would be driven by cost. We found that invasion and crop raiding primarily occur during the dry season and are associated with mangoes. Farmers are using many countermeasures. Effective countermeasures target giraffe behavior combined with physical barriers. Countermeasures are most effective when negative associations with humans are reinforced. Floodlights and speakers that play predator calls both reduce invasion, but only if used consistently. Overall, farmers’ perceptions matched giraffe dynamics. Availability was the most important factor in farmers’ willingness to try a countermeasure. Our results suggest that conflict can be reduced and there is interest from farmers in doing so, but use of countermeasures should be consistently applied and supported by making necessary equipment and instructions available.

## Background

Globally, humans are the leading cause of decline and extinction of large mammals, due to poaching, land-use conversion, and human-wildlife conflict (Ripple et al., 2017). Large mammals can be particularly prone to human-wildlife conflict because they require more space (Noonan et al., 2020), often roaming outside of formally protected areas where they are potentially snared, poached, or killed in retaliation for livestock or crop depredation (Acharya et al., 2016). The risk of conflict will likely increase in the future as previously contiguous wildlands become more fragmented (Zou et al., 2025), especially in areas where large mammals are found outside of protected areas (Bashir & Wanyonyi, 2024). Extreme declines in populations of many large mammals highlight the urgent need to improve the effectiveness of policies and strategies for large mammal conservation (Macdonald, 2019; Ogutu et al., 2016). Given the multiple interacting threats affecting large mammals, transdisciplinary research and approaches to support human–wildlife coexistence (Marchini et al. 2019), coadaptation (Carter & Linnell 2016), and implementable solutions are essential for addressing their conservation (König et al., 2020).

Crop damage by large mammals is one of the most commonly reported human-wildlife interactions outside protected areas and can have major impacts on conservation outcomes (Mukeka et al., 2019; Peterson et al., 2010). Crop raiding by wildlife—when animals eat, trample, or otherwise damage human crops—is influenced by multiple ecological factors, including activity timing, resource requirements, and competition (Gunn et al., 2014; Naha et al., 2020), as well as human choices, including the types of crops farmers grow and countermeasures they use (Bista & Song, 2022; Mukeka et al., 2019; Yogendra et al., 2024). Countermeasures are actions taken by people to prevent wildlife from accessing crops and can help to alleviate crop raiding (MusyoKi, 2014; Can-Hernández et al., 2019; Montgomery et al., 2022). It is important to understand what factors contribute to crop damage because of the negative effects of crop damage, including increased food insecurity, loss of livelihoods for people, and reduced support from locals for conservation (Kiffner et al., 2021; Tiller et al., 2021), and a potential increase in the use of lethal countermeasures against large mammals (Bulte & Rondeau, 2007; Tiller et al., 2021).

Unfortunately, we often lack understanding of how both ecological factors that drive animals and human choices can influence crop raiding, limiting our ability to effectively intervene. A lack of knowledge about what drives animals to raid crops makes targeted efforts to deter them difficult (Hill, 2018). Because weather patterns affect availability of forage, for example, crop raiding may be more prevalent during periods of drought (Calhoun et al., 2025), but this idea has not been widely tested. Crop choices by farmers are often climate- and market-driven, and therefore, may be predictable across regions (Nde et al., 2024), but decisions are also often driven by local knowledge or material availability (Montgomery et al., 2022). Furthermore, human perceptions of wildlife activity and countermeasure effectiveness are key factors in decisions that can impact effective conflict mitigation. When perceptions do not match wildlife activity, this can result in poor conservation outcomes (Kiffner et al., 2021). Understanding how farmers decide which conflict mitigation countermeasures to use, their perceived effectiveness, and reasons for their effectiveness is therefore essential for providing lasting solutions to crop raiding.

To address this, we examined crop raiding by reticulated giraffes (*Giraffa reticulata*) in Garissa County, Kenya. Reticulated giraffes in Garissa County provide a useful and important case study to examine crop raiding for several reasons. First, reticulated giraffes face significant conservation challenges—they are among the most recent large terrestrial mammals to be listed in the IUCN Red List of Threatened Species as Threatened with extinction, in part, because of human-wildlife conflict (O’connor et al., 2019; Coimbra et al., 2021). Second, Garissa County has arguably the among world’s largest reticulated giraffe population, but conflict between farmers and giraffes is becoming increasingly prevalent due to the area’s population growing larger and more sedentary as people in the area shift from pastoralist to agricultural and urban lifeways (Ali et al., 2026). Finally, because crop raiding by giraffes has not historically been common (Mukeka et al., 2019), it is poorly documented, and effective countermeasures and perceptions of these countermeasures are poorly understood (Gašparová et al., 2024).

Our research goal was, therefore, to understand ecological factors influencing giraffe activity and human decision-making and how those factors potentially interact. We identified four objectives to better understand the mediating factors that influence interactions of giraffes and farmers that ultimately influence crop raiding (Fig. 1). Our first objective was to understand how farm invasion by giraffes was influenced by local weather. Because wildlife movements often track dynamic resources (Abrahms et al., 2021), we hypothesized that farm invasion by giraffes and giraffe abundance in agricultural areas would be more prevalent during dry seasons than rainy seasons (Hypothesis 1). Our second objective was to understand how crop type influenced farm invasion. Because giraffes are selective feeders, we hypothesized that farms with the most palatable crops (e.g., mangoes) would be most likely to be invaded by giraffes (Hypothesis 2). Our third objective was to understand the perceived and experimental effectiveness of commonly used countermeasures. Measures should be in use if they are effective, so we assumed the general perception would be that countermeasures are effective (Hypothesis 3A). Because giraffes are tall and large, we hypothesized that reducing farm invasion using only physical barrier-based countermeasures (e.g., makeshift fences) was unlikely and that countermeasures that targeted giraffe behavior (e.g., causing or exploiting fear of humans) would be more effective (Hypothesis 3B). Finally, our fourth objective was to understand local farmers’ perceptions of interactions with giraffes and countermeasures. Specifically, we wanted to understand if perceptions of giraffe behavior by farmers matched giraffe behavior. We hypothesized that local perceptions would closely match documented giraffe activity (Hypothesis 4A). We were also interested in the factors that influence the adoption of new or altered countermeasures, as these are important for long-term uptake of effective countermeasures. Therefore, we were interested in perceptions of the success of countermeasures and factors influencing farmers’ willingness to change methods. We hypothesized that countermeasure adoption would be driven primarily by cost and availability (Hypotheses 4B).

**Figure 1:**
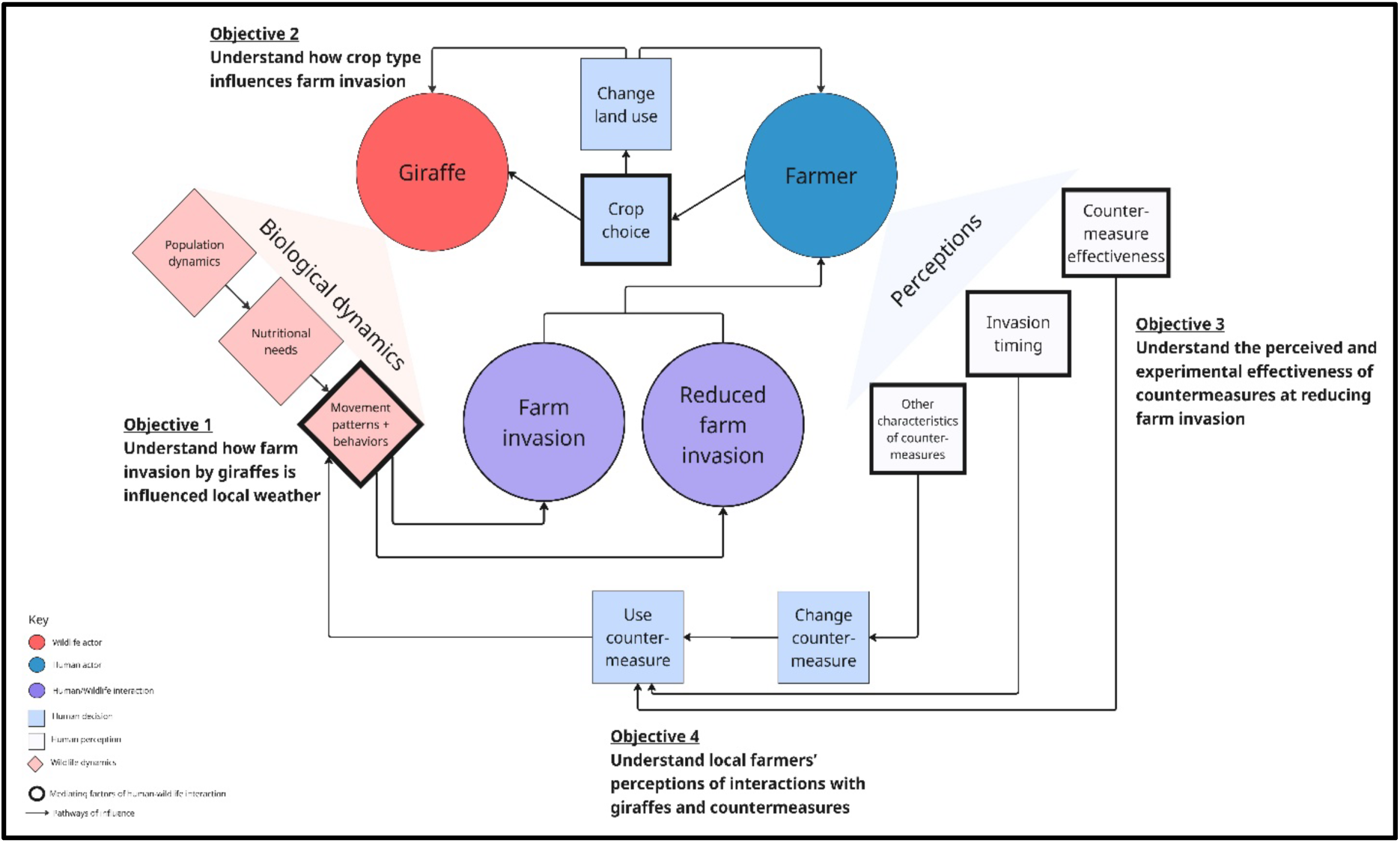
A conceptual framework showing the main mediating factors that influence giraffes (red circle) and farmers (blue circle) that result in invasion or reduced invasion (purple circles). Ecological factors that influence giraffes are shown in red (population dynamics, nutritional needs, and movement patterns and behaviors). Human perceptions (light blue) about invasion timing, countermeasure effectiveness and other characteristics of countermeasures are important precursors to decisions. Human decisions (medium blue) about using or changing countermeasures or crop choice can lead to changes in giraffe activity and ultimately result in interactions between giraffes and humans through farm invasion (purple). Each objective in this effort is designed to address a specific pathway of influence in this system, so it is mapped onto the framework accordingly.

## Methodology

### Study system

We conducted our research in 11 villages within Garissa Township, Garissa County, in eastern Kenya. This is also the location of Garissa Giraffe Sanctuary (hereafter “the Sanctuary”; 0°31′ S, 39°41′ E, 400m elevation), also locally known as Bour-algy Giraffe Sanctuary. The sanctuary was first conceptualized in 1995 and gazetted in 2000 for the protection of the reticulated giraffe (Ali et al., 2026). The sanctuary is managed by the Garissa County government and sits on approximately 125 km^2^ of land (Fig. 2). Although the sanctuary is a formally designated protected area, local pastoralists live within its boundaries with their livestock. The region is characterized by an arid to semi-arid climate, with rainfall ranging from 200 mm to 500 mm per year and falling in two wet seasons—March-May (long wet season) and October-December (short wet season). Temperatures are relatively high—maximum daily temperatures range between 28°C and 38°C, occasionally exceeding 40°C (Ali et al., 2017). The vegetation is composed of xeric shrubby and thorny bushes, and open savanna woodland dominated by *Acacia, Commiphora*, and *Combretum* species. *Acacia reficiens* and the invasive mesquite *Prosopis juliflora* are becoming increasingly abundant and causing large-scale shrub encroachment on this historically grassland ecosystem. Besides domestic livestock, the reticulated giraffe is the most abundant herbivore. Other common native species present include gerenuk (*Litocranius walleri*), Kirk’s dik-dik (*Madoqua kirkii*), lesser kudu (*Tragelaphus imberbis*), desert warthog (*Phacochoerus aethiopicus*), waterbuck (*Kobus ellipsiprymnus*), and spotted hyena (*Crocuta crocuta*). Livestock species present include goat (*Capra hircus*), sheep (*Ovis aries*), cattle (*Bos indicus*), camel (*Camelus dromedarius*), and donkey (*Equus asinus*). Most human residents are pastoralists of ethnic Somali descent. In the recent past, local residents have become more sedentary and practice farming along the Tana River, a major source of water in the region (Di Matteo, 2025).

**Figure 2:**
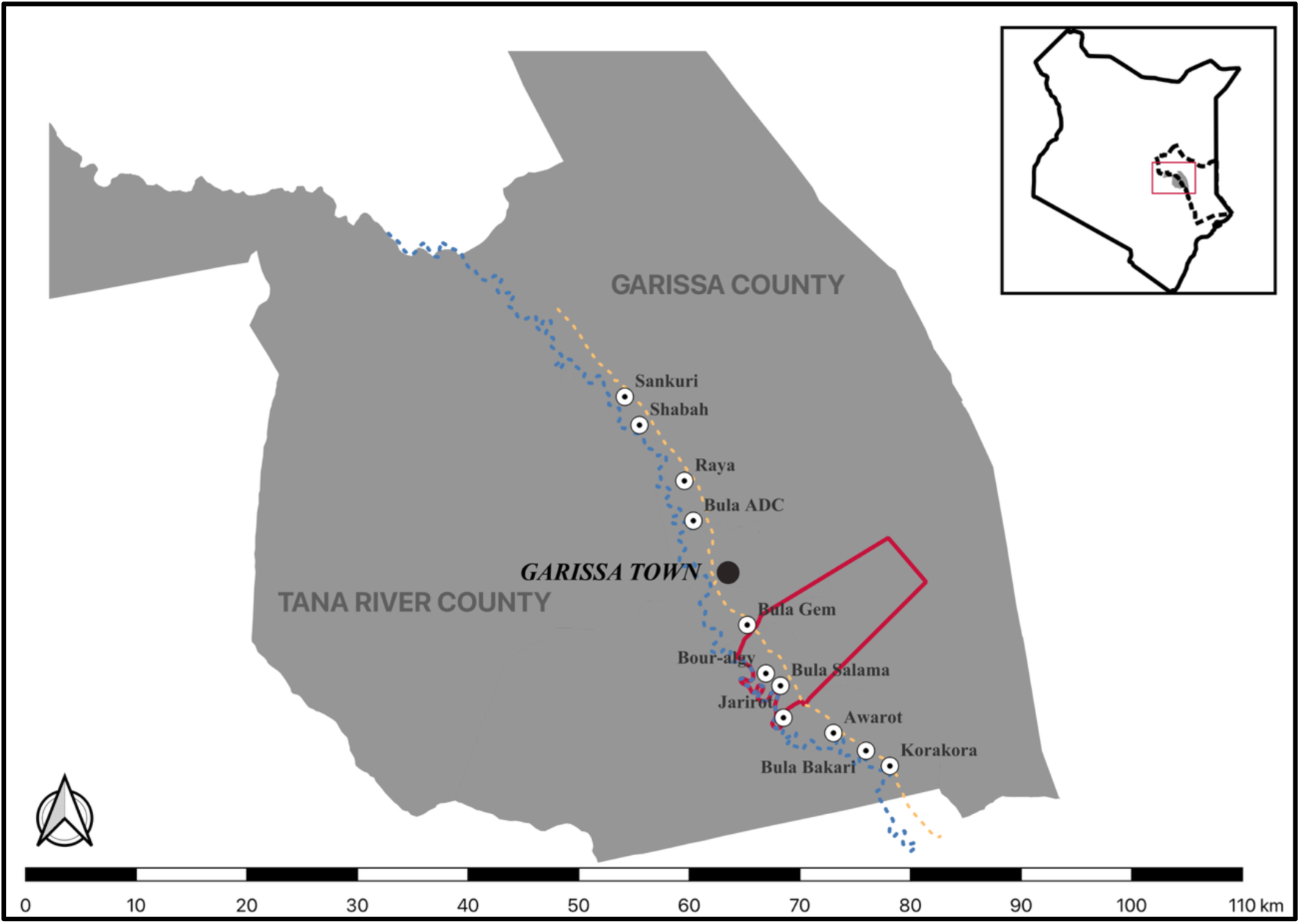
Map of our study site. The main map shows the Tana River (dotted blue line), the major source of water that divides Tana River (left) and Garissa (right) Counties. The red polygon represents the boundary of the Garissa Giraffe Sanctuary, while the dotted yellow line represents the major road in the region connecting counties to the north and south of Garissa. We used sections of this road as a transect for distance sampling and photographing giraffes. The white circles with black dots in the center mark the locations of the villages in which we administered our questionnaires. Insert map shows the boundary of Kenya (black solid line), Garissa County (black dotted line), and our study site (grey area in red rectangle).

### Data collection

We drew from three sources for data collection: direct observation of giraffes; survey data from local farmers; and experimental implementation of countermeasures by local farmers. Each source involved different data collection methods, described below.

### Giraffe observations

To examine how farm invasion by giraffes was influenced by local weather (Objective 1), we gathered precipitation data and estimated local giraffe density and abundance across a section of short-wet (December) and dry (January to May) seasons. We obtained values for precipitation from Climate Hazards Center InfraRed Precipitation with Station data accessed via Google Earth Engine (Funk et al., 2015; Gorelick et al., 2017). Because of the lag between precipitation and reagreeing of vegetation (Godlee et al., 2024), and the sparse nature of rainfall in Garissa, we used pentad image collection. To collect data on giraffe density and abundance, we first identified a road (dotted yellow line in; Fig. 2) that splits the sanctuary into two unequal parts, (1) the riverfront area where farms are located along the Tana River, and (2) upland areas where, except for scattered settlements, there are no farms. Giraffes were reported by locals to cross this road toward the Tana riverfront late in the evening (supposedly to invade farms at night) and back toward the uplands in the morning to avoid human activity in the farms. We used two survey types: line transect surveys and photographic mark-recapture surveys.

To estimate giraffe density using line transect distance sampling (Buckland et al., 2015), we conducted transect surveys using a motorcycle (Bajaj Boxer BM-100, Pune, India) moving at a speed of 10km/hr with two observers—one looking on each side of the road. We used a rangefinder (Nikon 8397 ACULON AL11, Tokyo, Japan) to estimate the sighting distance and a compass (Suunto A-10, Vantaa, Finland) to measure the transect and giraffe bearings, which were then used to calculate sighting angles. We conducted transect surveys for 19 days (4 in December 2021, 3 in January, 5 in March, 3 in April, and 4 in May 2022) using established roads segmented to variable lengths within the study sites (Augustine, 2010). We sampled Bula Gem and Bour-algy together (3.1 km), but Bula Salama (3.1km), Jarirot (3.2km), and Awarot (5.7 km) as separate segments. The final data was pooled since these segments were not biologically significant but arbitrary boundaries marked by large luggas (i.e., intermittent washes) and named after the nearest village center. We collected data early in the morning (0600 to 0800hrs) and late in the evening (1700 to 1900hrs) when giraffes were reportedly seen moving away from and towards the Tana River to avoid human activities or visit farms, respectively. Because temperature in Garissa is often high during the day, this also coincided with time giraffe were most active.

Additionally, to estimate the number of giraffes that might be involved in farmer-giraffe conflict, we identified individuals through photography and used mark-recapture to estimate abundance from these photographs (Bolger et al., 2012; Halloran et al., 2015). Using a hand-held Digital Single-Lens Reflex camera (Cannon EOS Rebel *T3i*, Canon Inc., Tokyo, Japan) we took photos of giraffes for 5 days in January, 1 day in February, 16 days in March, and 2 days each in April and May 2022. We photographed the whole body of each giraffe from the left flank and uploaded it to the GiraffeSpotter web platform (Berger-Wolf et al., 2017) for individual identification (Lee et al., 2022). We assigned the giraffes’ sexes and approximated age based on observable attributes, such as the presence of observable genitalia or the nature of ossicones (i.e., giraffes’ hornlike structures).

### Survey data from farmers

In 2022, between January and July in 11 villages within Garissa Township, the first author and a translator verbally administered 207 semi-structured survey each with 60 questions to local farmers in the study area (see Supplementary Material 1). The survey was designed to determine the perceptions of farmers concerning farm invasion by giraffes (Objectives 2 and 4). We used a snowball sampling approach by asking participants to recommend other potential participants (hereafter, farmers). With the survey, we collected information about the demographics of the farmers, information about their farms, perceptions about giraffe invasions, information and perceptions about countermeasures and their effectiveness, and attitudes about trying new countermeasures and which countermeasures they were willing to try. The questionnaire was primarily administered in Kiswahili, but it was administered in Somali when farmers were not comfortable communicating in Kiswahili. All survey responses were recorded electronically using a mobile application (ODK Collect, Seattle, US). Because the survey was originally intended to inform the first author’s on-the-ground conservation work rather than research, he did not seek Institutional Review Board approval for this survey. After beginning his Ph.D., however, the authors submitted this survey for retroactive reviewed by the University of Arizona Institutional Review Board, who deemed it “not human research” and approved interview protocol and sampling approach after a partial review (permit #STUDY00006143).

### Experimental implementation of countermeasures by farmers

To understand the potential effectiveness of different countermeasures (Objective 3), we experimentally tested the use of solar-powered motion sensor floodlights and predator calls (i.e., a lion roaring) to reduce farm invasion by giraffes. We used the 180W solar monitoring sensor lamp (CL 877B, China) with motion sensor range of 5-10m and set to only activate when an object passed by. We noticed that the motion sensor range was most effective within the 5m mark, we therefore, mounted the solar floodlights on 5m tall posts spaced 9m apart to ensure any giraffes moving between any two posts could activate a floodlight. We obtained a lion roar from a local ranger (S. Haji, personal communication), trimmed it to 30 seconds and added 4.5, 9.5, and 14.5 minutes each of blank recording to make the sound cycle through at five, ten, and fifteen-minute intervals, and saved it to memory cards. We issued study participants solar speakers (Golon X-BASS RX-BT190S, Quanzhou, China) with the three sounds programmed to play on a random shuffle in a loop. We selected 72 farms based on willingness to participate and assigned each of the selected farms to one of four experimental groups: (1) light control (no lights provided), (2) sound control (no sound provided), (3) light treatment (lights installed on posts), and (4) sound treatment (solar speaker with roars on loop). We then gave farmers at each farm a printout calendar to mark days on which giraffes invaded the farm. This research was covered by Kenya Wildlife Research and Training Institute permit number WRTI-0119-11-21. Because the first author conducted fieldwork independently and was based at an institution that does not have an Animal Care and Use Committee, this research was not approved by an Animal Care and Use Committee, but it followed the 2016 Guidelines of the American Society of Mammalogists for the use of wild mammals in research and education (Sikes & the Animal Care and Use Committee of the American Society of Mammalogists, 2016).

### Data analysis

We performed all statistical analyses in the R statistical software environment (version 4.3.1; R Core Team, 2023).

### Giraffe observations

We modeled the relationship between the observed number of giraffes and average precipitation each day using a Generalized Additive Model (GAM) to account for nonlinear associations. We then estimated detection probability and density of giraffes using hierarchical distance sampling model in the ‘unmarked’ R package (version 1.4.3; Kellner et al., 2023). Because the farthest perpendicular distance observed was 503m and because giraffes are large and can be seen from distances greater than 1000m in relatively open areas like Garissa, we did not truncate observation distances. We split the data into six bins of 100m each to cover all perpendicular distances. After visually inspecting the distance histogram, we fit four candidate models with hazard-rate detection functions including, (i) a null model with both detection and density held constant, (ii) detection varying with precipitation, (iii) density varying with precipitation and (iv) both detection and density varying with precipitation. We used AIC to select the best model (Akaike, 2003).

To estimate the number of giraffes in our study areas, we pooled the 26 survey days by the months January (Days 1–5), February (Day 6), March (Days 7–22), April (Days 23–24), and May (Days 25–26). This represented different sampling efforts for each sampling session. Because of the relatively short sampling periods, we assumed both demographic and spatial closure and fitted a closed population capture-recapture model using the ‘Rcapture’ package (version 1.4.4; Rivest & Baillargeon, 2007) and used AIC to select the best model. We minimized model complexity and accounted for sparse capture history by pooling capture occasions into 5 occasions for each month. Pooling not only reduced the number of capture occasions but also facilitated model convergence and reliable estimation of abundance. To account for varying sampling effort, we implemented a Bayesian closed population model in ‘rjags’ (version 4.17; Plummer, 2008) and modeled detection probability as a logistic function. We augmented the observed detection matrix to 1000 individuals and used uninformative priors for the intercept (α∼ N(0,0.01)), effort effect (β∼ N(0,0.01)), and inclusion probability (ψ∼ uni(0,1)). We computed posterior summaries and convergence diagnostics using the ‘coda’ R package (version 0.19.4.1; Plummer et al., 1999) with three chains of 5,000 iterations and 100 burn-in iterations ensuring the model converged with all R^ near 1.0 (Brooks & Gelman, 1998).

### Survey data from farmers

We used a series of steps to clean and analyze the data obtained from our survey of farmers. To clean the data, we first visually checked for any field entry errors. Next, we recoded responses into binary variables where appropriate or to split multi-faceted responses into separate columns. For example, for the question “Have you experienced giraffe invasion on your farm?” we coded yes and no responses into 1 (yes) and 0 (no). For the question “What crops do you grow?” we coded responses for mangoes, vegetables, citrus, and melons into individual columns of 1 (farmer grows that crop) or 0 (farmer does not grow that crop).

To understand how crop type influenced farm invasion (Objective 2), we used R/RStudio to conduct two-tailed Fisher’s exact tests to determine if giraffe invasion was related to the presence of palatable crops (mangoes) or other crops grown. We used Fisher’s exact test to account for the small sample sizes (1-5 farms) across some of the groups.

### Experimental implementation of countermeasures by farmers

To determine the effectiveness of countermeasures (Objective 3), we estimated the deterrence success rate of the solar floodlight and sound by dividing the total number of days farmers reported the method working by the number of days the experiment ran on each farm. We performed a non-parametric Wilcoxon rank-sum test to compare farmer-recorded deterrence success rates between light and sound treatments and controls.

To understand farmers’ perceptions of giraffe activity and countermeasures (Objective 4) we examined both categorical and free response questions. For free response questions that related to our objectives, we determined appropriate code categories and coded responses according to these categories. Surveys focused on which countermeasures farmers used to mitigate giraffe-farmer conflict, and farmers were also asked to provide a reason as to why they thought a countermeasure was effective or not. Although the responses were diverse, they revealed that farmers conceptualized the effectiveness or ineffectiveness of countermeasures resulting from four main challenges. When dealing with problem giraffes, farmers tried to (1) physically prevent them from accessing resources by reducing the advantages of giraffe anatomy (particularly height and size; hereafter “anatomy”), and (2) take advantage of animal behaviors like fear of stimuli (“behavior”). However, (3) giraffes often learned over time that countermeasures did not have negative outcomes, which decreased their efficacy (“learning”), and (4) environmental factors could interfere with proper implementation of a countermeasure (e.g., some farmers reported that presence of hippopotamuses made the use of human guards dangerous, while others reported that fire was a potential hazard to humans and property; “environment”). For free response questions, we coded the farmers’ responses into these four categories.

## Results

Results are presented below by objective. Table 1 provides an overview of our main results organized by research objectives, including our hypotheses, data collection and analysis methods, and the table or figure where those results can be examined in detail.

**Table 1.** An explanation of our main research objectives, hypothesis, methods, main results, and table or figure aligned with those results.

| Objective | Hypothesis | Method | Result | Table or Figure |
| --- | --- | --- | --- | --- |
| 1) Understand how farm invasion by giraffes is influenced local weather | H1: Farm invasion by giraffes and giraffe abundance in agricultural areas would be more prevalent during dry seasons than rainy seasons | Data collection: Giraffe observations<br>Data analysis: GAM | Lower giraffe abundance with higher rainfall | Fig. 3 |
| 2) Understand how crop type influences farm invasion | H2: The presence of palatable crops (e.g., mangoes) drives farm invasion by giraffes | Data collection: Survey data from farmers<br>Data analysis: Fisher's exact test | Presence of palatable crops (mangoes) is significantly associated with giraffe invasion, whether farmers grow multiple crops ($p = 0.047$) or just single crops ($p = 0.003$) | Table 1 |
| 3) Understand the perceived and experimental effectiveness of commonly used countermeasures at reducing farm invasion | H3A: Most countermeasures would be perceived as effective | Data collection: Survey data from farmers<br>Data analysis: Descriptive statistics | Most people experienced invasion. Those who did not used guards, fences, or fire (Fig. 5C)<br>Most people were neutral or positive about countermeasures, except for fences (most people were negative about fences; Fig. 6)<br>Ineffectiveness of fences was largely attributed to giraffe anatomy | Fig. 5C<br>Fig. 6 |
|  | H3B: Experimentally reducing farm invasion using only physical barrier-based countermeasures | Data collection: Experimental implementation of countermeasures by farmers | Applying a countermeasure (light or sound) resulted in a significant reduction in farm invasion by | Fig. 8 |
| | (e.g., makeshift fences) was unlikely and that countermeasures that targeted giraffe behavior (e.g., causing or exploiting fear of humans) would be more effective | Data analysis: Wilcoxon rank-sum test | giraffes ($W = 289, p < 0.0001$).<br>Use of sound devices (i.e., predator call) resulted in a significantly higher deterrence rate ($W = 100, p\text{-value} = 0.015$) than using light. | |
| 4) Understand local farmers' perceptions of interactions with giraffes and countermeasures<br>a) Understand if perception of giraffe behavior by farmers matched with our measurement of giraffe activity | H4A: Local perceptions would closely match documented giraffe activity | Data collection: Survey data from farmers<br>Data analysis: Descriptive statistics | Most people experienced invasion only in the dry season | Fig. 4A |
| b) Understand the factors that influence the adoption of new or altered countermeasures | H4B: Countermeasure adoption would be driven primarily by cost and availability | Data collection: Survey data from farmers<br>Data analysis: Descriptive statistics | Most people were extremely interested in trying new countermeasures, and the most common reason they cited was ease of use, followed by availability, neighbor use, Kenya Wildlife Service use, and affordability (Fig. 7)<br>When asked what they were willing to try/improve, most people said fences, followed by lights, guard, unsure, and financial support | Fig. 7 |

### Objective 1: Effects of precipitation on the density and abundance of giraffe within farmland

Using distance sampling, we estimated the density of giraffes in our study site to be about 6 giraffes/km^2^ (5.9 giraffes, 95% CI: 2.9 – 12.2), or 737 giraffes in our 125km^2^ study area. Using photographic mark-recapture, we estimated the number of giraffes within our study area during that period to be about 603 (95% CI: 425 – 780) giraffes and 669 (95% CrI: 476 – 914) giraffes, using a model in which capture probability varied with time (from the ‘Rcapture’ package) and one that accounted for varying sampling efforts (the Bayesian model), respectively.

We estimated fewer giraffes as rainfall increased within our study area (Fig. 3). When there was no rainfall (0mm), we estimated about 34 giraffes (34.27 giraffes, 95% CI: 19.19 – 49.35) per day. When rainfall increased above 10mm per day, we expected to count about 5 giraffes per day (5.17 giraffes, 95% CI: −6.32 – 16.65), but their counts were highly uncertain. Similarly, out of 194 farmers that reported having experienced farm invasion by giraffes, a majority (178; 92%) of the farmers indicated that farm invasion by giraffes, occurred mostly during the dry season (coinciding with late January through early May; Fig. 4A).

**Figure 3:**
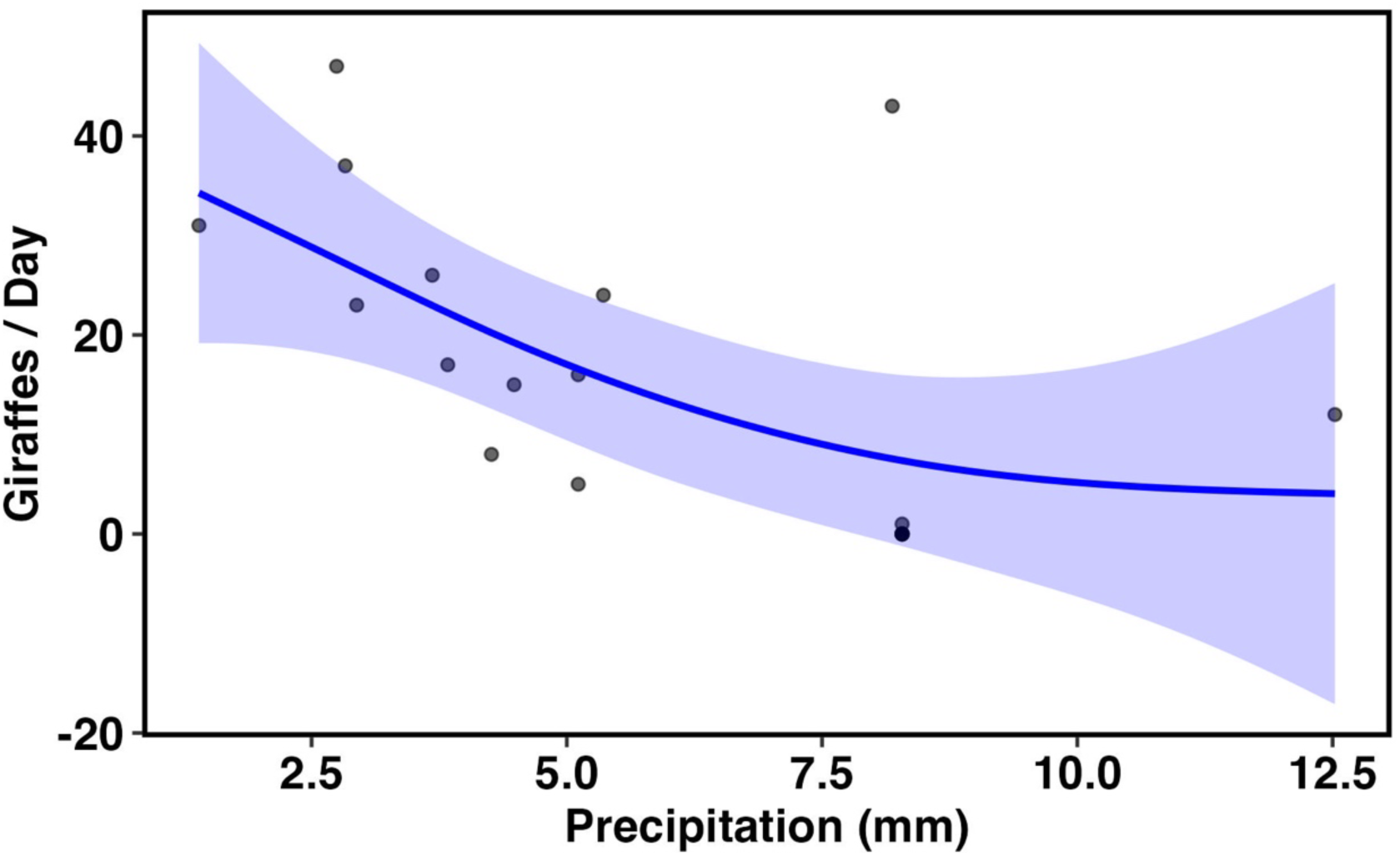
Scatterplot illustrating the total number of giraffes sighted each day as a function of daily precipitation. The blue line is the modeled relationship showing that as precipitation increases, the number of giraffes sighted each day decreases.

**Figure 4:**
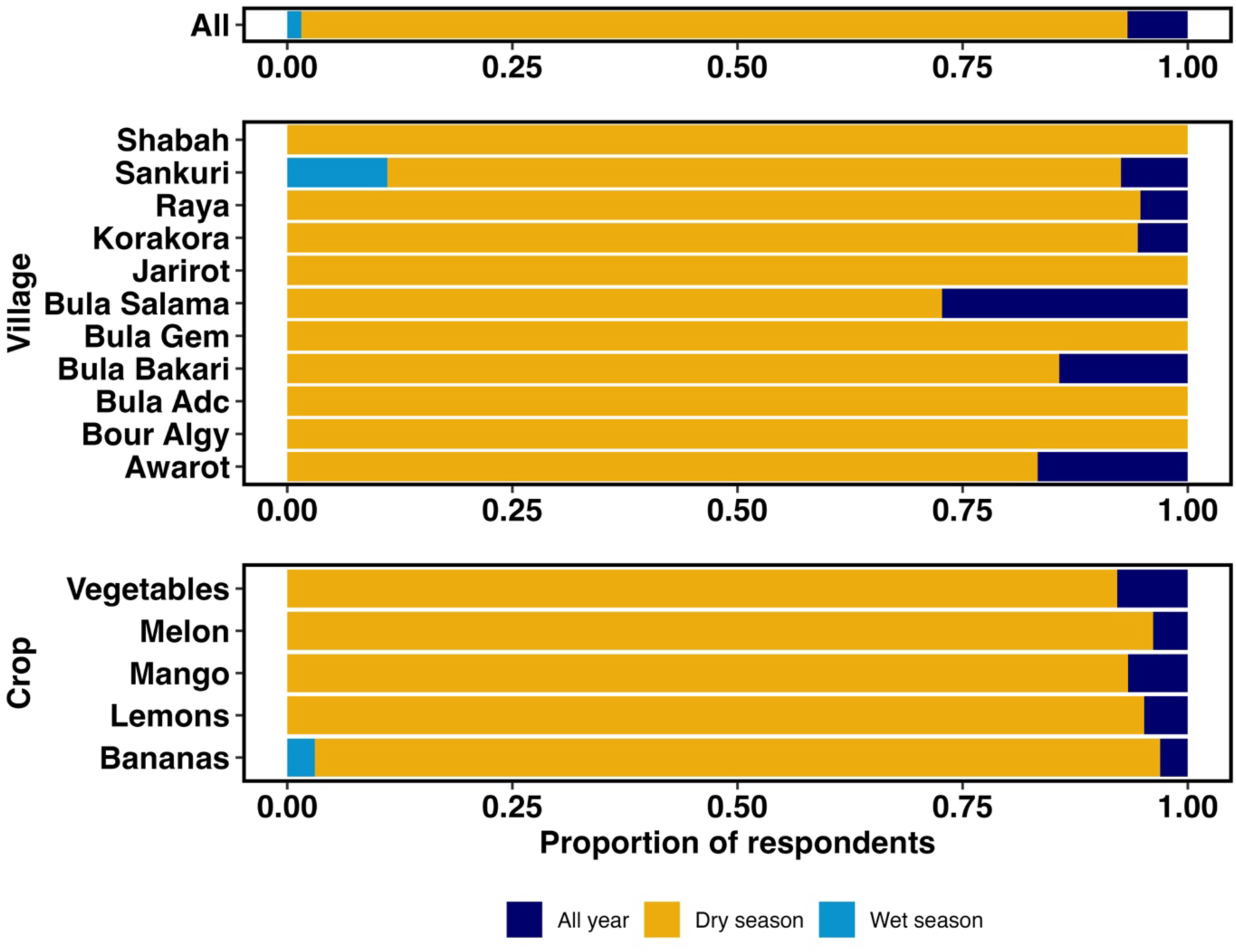
Stacked bar chart illustrating (A) the proportion of farmers who thought giraffe invasion occurred all year (dark blue), only in the dry season (light blue), or only in the wet season (light blue); (B) the proportion of farmers by village who thought giraffe invasion occurred all year, only in the dry season, or only in the wet season; and (C) the proportion of farmers by the type of crop grown who thought giraffe invasion occurred all year, only in the dry season, or only in the wet season.

### Survey data from farmers

#### Locations and demographics

In total, the first author administered 207 surveys, each of which lasted approximately 30 to 60 minutes. Interviews were distributed as follows in the 11 villages: Awarot (*n* = 18); Bour-algy (*n* = 10); Bula ADC (*n* = 18); Bula Bakari (*n* = 21); Bula Gem (*n* = 24); Bula Salama (*n* = 11); Jarirot (*n* = 17); Korakora (n = 20); Raya (*n* = 19); Sankuri (*n* = 38); Shabah (*n* = 11); Fig. 2). Many of the farmers were male (152 individuals; 73%). Their age, however, was distributed relatively evenly, with the largest categories represented by groups between the ages of 29 and 39 (45 individuals; 22%), 40 and 49 (47 individuals; 23%), and 50 and 59 years old (51 individuals; 25%).

Most farmers (119 individuals; 58%) reported having only attained informal education (i.e., they had only attended *madrassas*–religious classes to recite and learn the Quran). Most farms (149 farms; 72%) were small (i.e., less than 5 hectares). Farmers grew a variety of crops on their farms, with most farmers (169 farms; 82%) growing two or more crops. Mangoes (182 farms; 88%), citrus fruits (147 farms; 71%), bananas (102 farms; 49%), vegetables (57 farms; 28%), and melons (55 farms; 27%), respectively, were the most frequently grown crops.

### Objective 2: Relationship of crop type to farm invasion

Most farmers (194 individuals; 94% of all farmers surveyed) reported that giraffe invasion had occurred on their farm (Fig. 5A). The small proportion that reported that invasion had not occurred (13 individuals: 6%) were in two villages: Sankuri (11 individuals) and Korakora (2 individuals; Fig. 5B). All farmers in our survey grew crops, but the type and amount varied. Most farmers grew two or more crops (169 farms; 82% of farms sampled). For those that grew two or more crops, giraffe invasion was significantly related to growing mangoes (*p* = 0.047). This relationship was also significant for the farms who only grew single crops (38 farms; 18% of farms sampled) (*p* = 0.003; Table 2).

**Figure 5:**
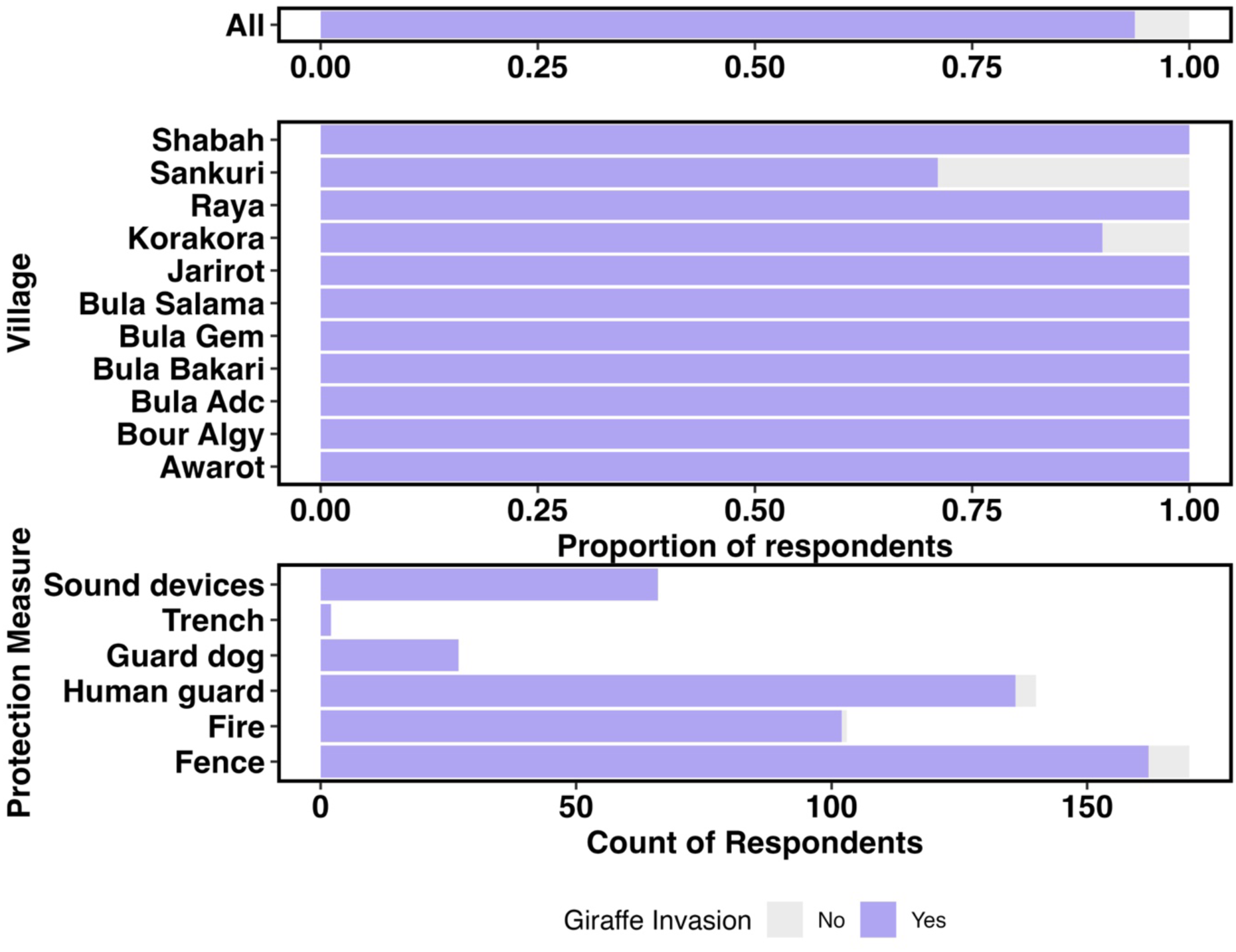
Stacked bar charts illustrating: (A) the proportion of farmers who did (purple) or did not (gray) experience giraffe invasion on their farm; (B) the proportion of farmers by village who did (orange) or did not (blue) experience giraffe invasion on their farm; and (C) the count of measures used by individuals who did (orange) or did not (blue) experience giraffe invasion. Some individuals used multiple measures, so counts add up to more than the 207 farmers.

**Table 2.** Contingency tables for the 169 farms that grew multiple crops (top) and the 38 farms that grew only a single crop (bottom) as well as the associated p-values for the two-sided Fisher’s exact test.

| | Farms not growing mangoes | Farms growing mangoes | Fisher's exact test $p$-value |
| --- | --- | --- | --- |
| Giraffe invasion | 3 | 164 | 0.047 |
| No giraffe invasion | 1 | 1 |  |
| | Single crop farm not growing mangoes | Single crop farms growing mangoes | Fisher's exact test $p$-value |
| Giraffe invasion | 10 | 17 | 0.003 |
| No giraffe invasion | 10 | 1 |  |

### Objective 3: Understand the perceived and experimental effectiveness of countermeasures

All farmers (194 individuals; 94% of respondents) that had experienced farm invasion used more than one countermeasure to reduce farm invasion by giraffes. Across all countermeasures that individual farmers used (e.g., guards, fences, fire, guard dogs, trenches, and sound-producing devices; Supplementary 1, Figs. S1-S4), many farms experienced invasion (Fig. 5C). On the 13 farms that did not experience invasion, countermeasures used included guards, fences, and fire, plus one farm with no countermeasures (Fig. 5C). Of the 194 farms that had experienced invasion all used countermeasures, most used fences (157 farms; 81%), followed by human guards (138 farms; 71%), fire (99 farms; 51%), sound producing devices (64 farms; 32%), flashlights (34 farms; 18%), guard dogs (27 farms; 14%), trenches (2 farms; 1%) and scare crows (1 farm; 0.5%). Note that because some individuals used multiple countermeasures on their farms, the totals add up to more than 194 and percentages reflect the percentage of respondents within the invasion group that had used that countermeasure.

Perceptions of the effectiveness of these countermeasures among the 194 farmers varied. Because individuals within this group used many different countermeasures and sometimes multiple countermeasures, the percentages presented in the following two paragraphs reflect the proportion of individuals who used that countermeasure, a subset of the 194 total. Fences were perceived as a largely ineffective countermeasure, with a majority of farmers reporting fences to be ineffective (111 farmers; 71%) compared to fewer farmers (8 farmers; 5%) reporting them to be effective (Fig. 6). Human guards were perceived as effective, with many farmers (62 farmers; 39%) reporting human guards to be effective compared to only a few farmers (4 farmers; 2.5%) reporting them to be ineffective. Fire was perceived as an ineffective countermeasure, with many farmers (47 farmers; 47%) reporting fires to be ineffective compared to few farmers (4 farmers; 4%) reporting that fire is effective. Sound-producing devices were perceived to be of mixed effectiveness, some farmers (14 farmers; 22%) reporting they were effective but a similar number of farmers (9 farmers; 14%) reporting them to be ineffective. Flashlights were also perceived as being of mixed effectiveness with some farmers (8 farmers; 24%) reporting they were effective but a similar number of farmers (10 farmers; 29%) reporting them to be ineffective. Guard dogs were also perceived as ineffective, with only a single farmer (1 farmer; 4%) reporting they were effective, compared to several farmers (9 farmers; 33%) who reported them to be ineffective. Only three farmers reported using trenches and scarecrows, but all three reported that these countermeasures were effective.

**Figure 6:**
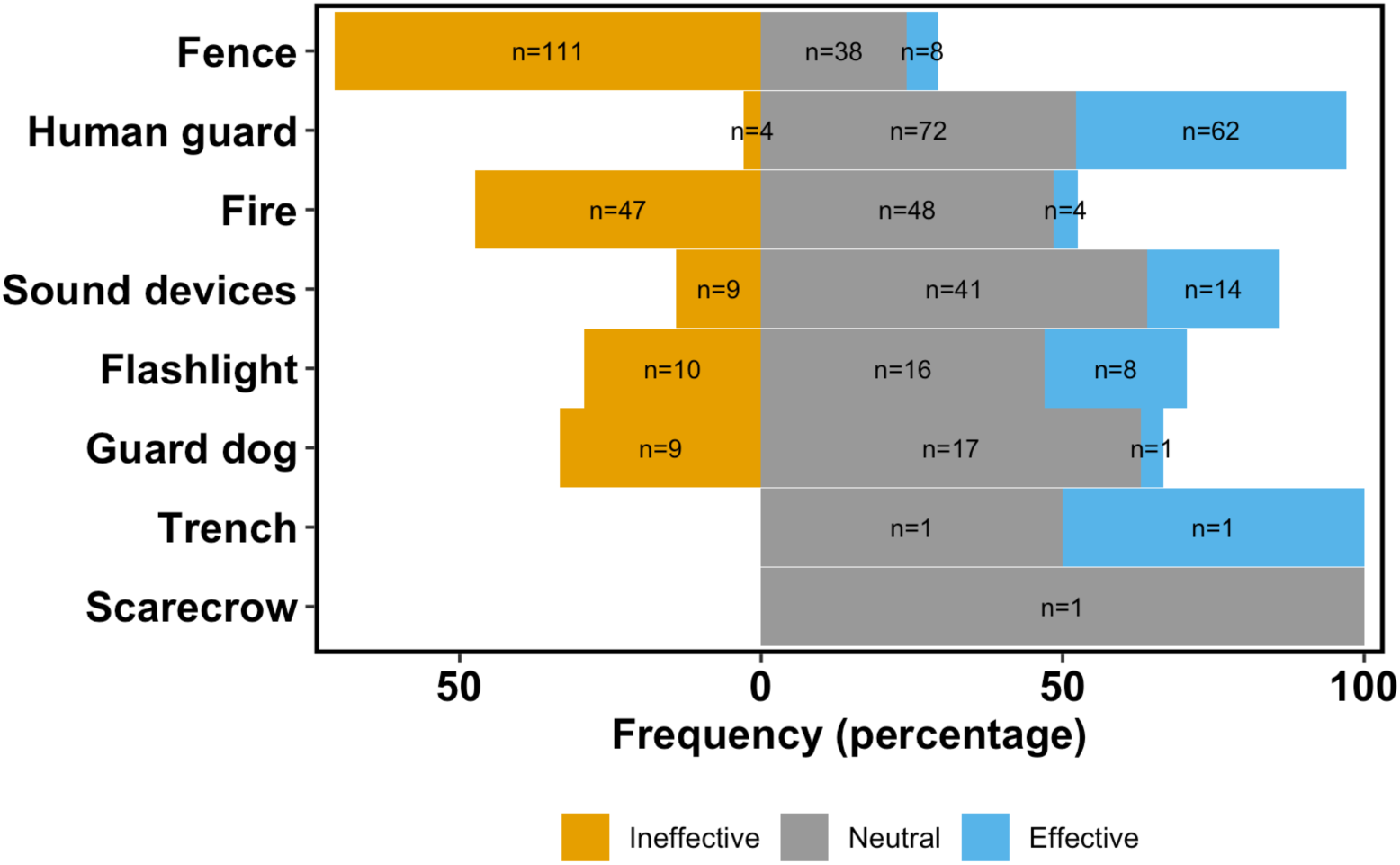
The percentage (bar length) and total number of farmers (text on bars) who mentioned the efficacy of countermeasures and whether or not they thought countermeasures were ineffective (negative; orange), neutral (gray), or effective (positive; blue).

The ineffectiveness of fences was largely attributed to giraffe anatomy. Many farmers (110 individuals; 76%) reported that giraffes walked over, broke, or went around fences. Effectiveness of human guards was attributed to the ability of humans to manipulate giraffe behavior, and ineffectiveness arose from environment—when effective, human guards could impact giraffe behavior by chasing them from the farm, but when ineffective, it was due to late nighttime invasions (when guards fell asleep) or safety concerns from hippos. Fire was considered ineffective because of learning—without additional measures (e.g., human guards), giraffes learned that fires alone pose no risk to giraffes, and some (4) farmers were concerned about environmental factors (e.g., risk of fires spreading on the farm). The effectiveness of sound-producing devices was attributed to behavior (giraffes associated sound-producing devices with human guards who rattle iron sheets or shake bottle with rocks insides when chasing giraffes from farms). There were, however, reports of sound-producing devices becoming ineffective due to learning when human guards were not used in conjunction with sound-producing devices. Similarly, farmers believed flashlights were effective when giraffes associated them with human guards who used flashlights when chasing giraffes from farms, but learning rendered the method less effective when not used in conjunction with human guards. As a results, farmers reported that flashlights were equally effective as they were ineffective in affecting giraffes’ behavior. Finally, guard dogs were reported to be effective due to behavior only when used in conjunction with human guards.

At 72 different farms, we assigned 34 farms to controls (18 light, 16 sound) and 38 farms to treatments (20 light, 18 sound). These experimental results indicated that applying a countermeasure (light or sound) resulted in a significant reduction in farm invasion by giraffes (W = 289, *p* < 0.0001). Furthermore, the use of sound devices (i.e., predator call) resulted in a significantly higher deterrence rate (*W* = 100, *p-*value = 0.015) than using light.

### Objective 4: Understand farmers’ perceptions of interactions with giraffes and willingness to try new countermeasures

Most farmers surveyed said that giraffe invasion occurred during the dry season (Fig. 4A). Out of 194 farmers that had reported farm invasions by giraffes, many (178 farmers; 92%) reported farm invasion to have occurred only during dry season, while the remaining few (13 and 3farmers; 6% and 2%) reported farm invasion have occurred year-round and only in the wet season, respectively.

Most farmers (132 farmers; 64%) were extremely interested in trying new countermeasures, but some were slightly (38 farmers; 18%), moderately (13 farmers; 6%) and not interested (24 farmers; 11%). For those who were extremely interested in trying new countermeasures (132 farmers; 64%), each farmer selected more than one reason why they were willing to try a countermeasure. The most important factors for willingness to try new countermeasures were local availability (90 farmers; 68%), followed by ease of use (76 farmers; 57%), use by trusted contacts like neighbors (67 farmers; 51%). Use by government officials (i.e., Kenya Wildlife Service) and cost were important for far fewer farmers (4 and 6; 3% and 4%, respectively; Fig. 7). Furthermore, we also asked farms to list a countermeasure they were willing to try. Out of 131 that responded, most farmers (95 farmers; 59%) suggested using variations of fences (e.g., electric, wires, wall etc.), while few suggest using light (16 farmers; 12%), human guard (15 farmers; 11%), and the government (5farmers; 4%) would help reduce farm invasion by giraffes respectively. Only one person wanted to try digging a trench.

**Figure 7:**
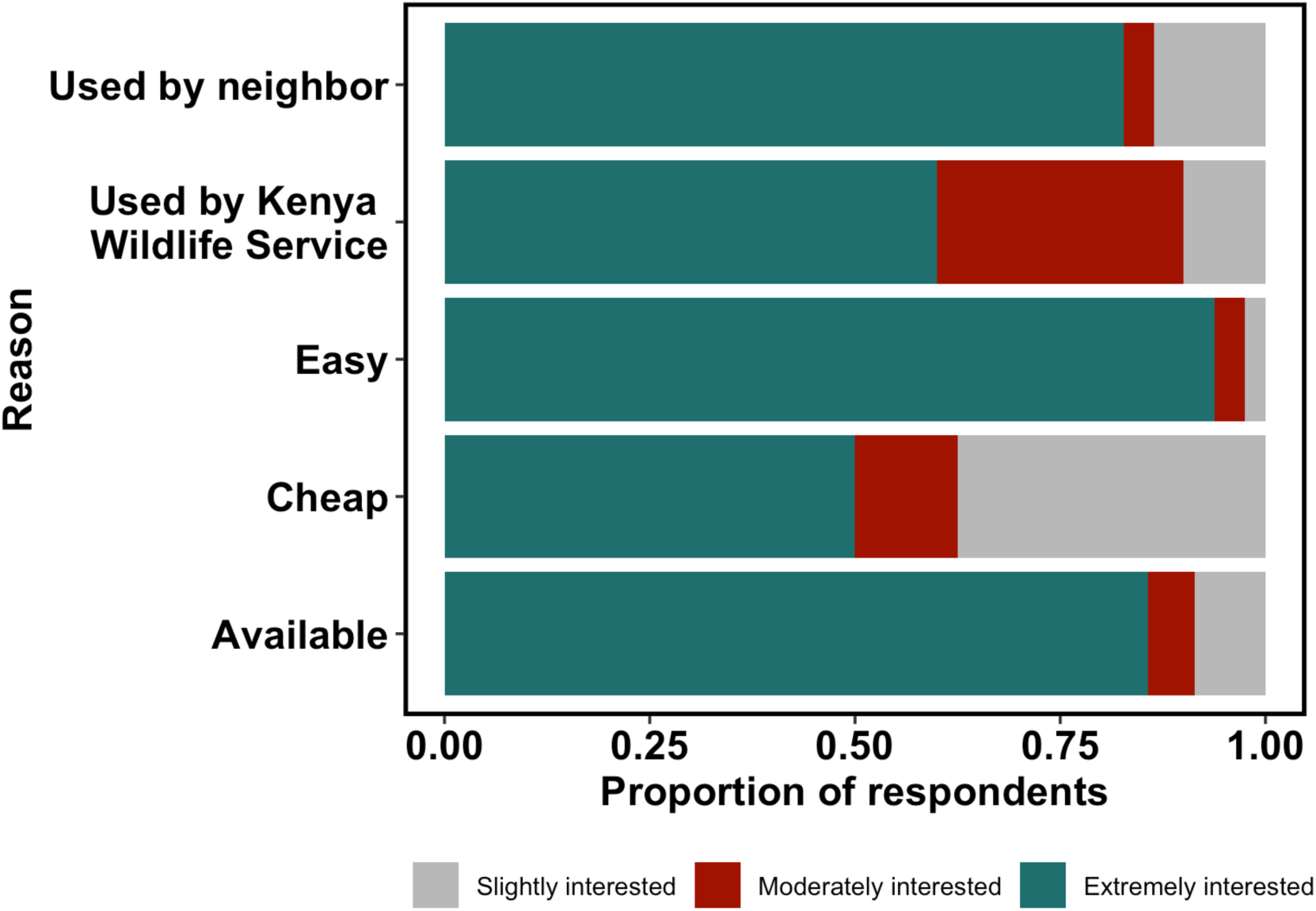
A graph showing factors influencing farmers decisions to trying a new method of reducing farm invasion by giraffes. The colors teal, red and gray represent farmers that responded they were extremely, moderately and slightly interested in trying a new method.

**Figure 8:**
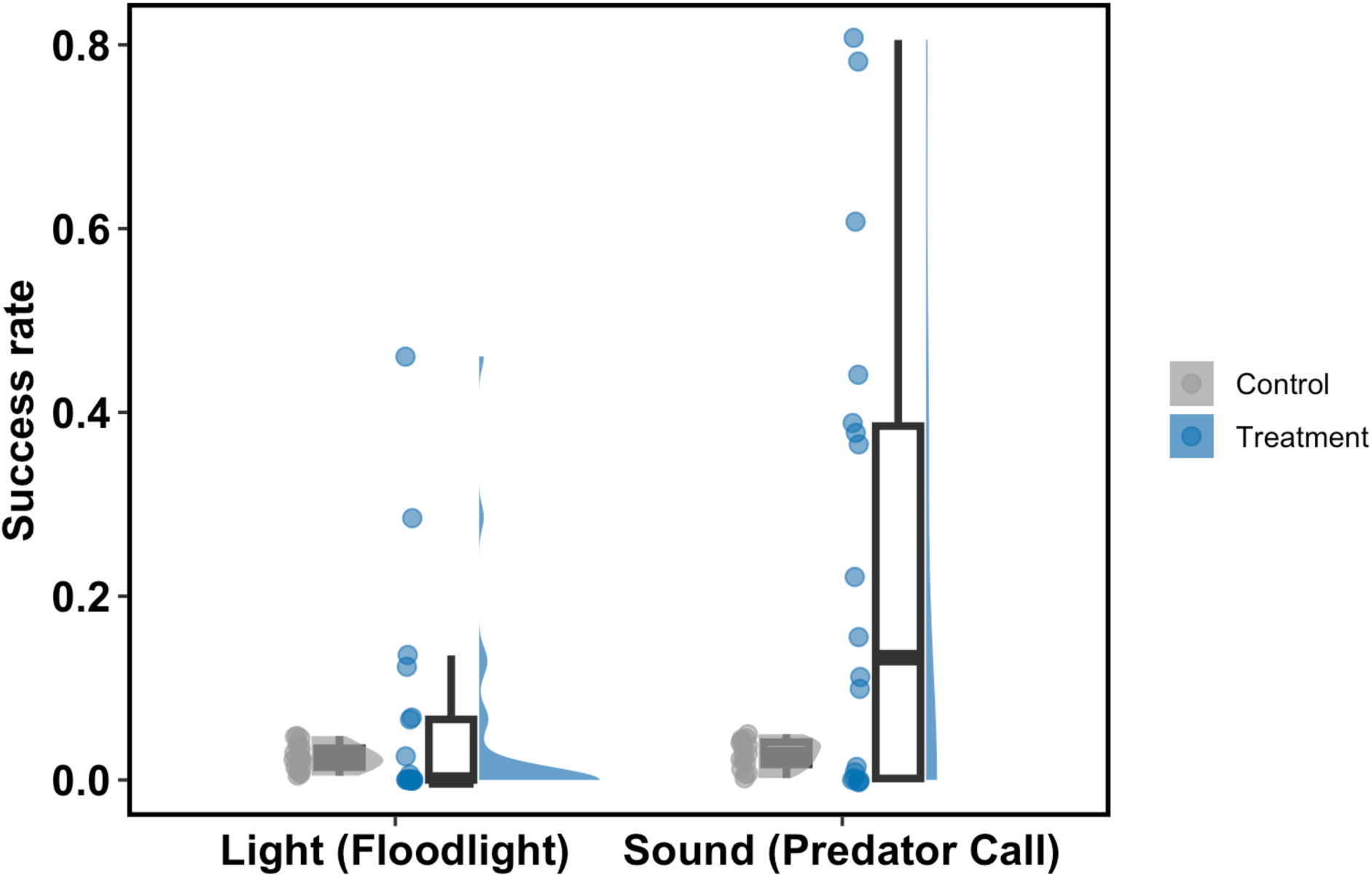
A graph showing the success rates (proportion) of solar-powered motion sensor floodlights (red) and predator calls broadcasted via Bluetooth speaker. The gray dots and associated boxplot represent control while blue dots, associated box plot and density plot represents treatment for each category. Because the farmers in control group reported daily invasion, the success rate was therefore zero but during plotting we added random value with maximum value of 0.05 for control to show up on the plot.

## Discussion

Better understanding the nature of human-wildlife conflict can help conservation practitioners develop better tools to mitigate it. In Garissa, conflict between farmers and giraffes primarily occurs during the dry season and appears to be driven in large part by the production of mangoes. Farmers are attempting many mitigation measures (often in conjunction), but the most effective measures target giraffe behavior rather than anatomy, and are most effectively deployed when paired with human guards who reinforce negative associations with humans and prevent giraffes from learning that countermeasures have no real negative impacts over time. These findings are similar to other human-wildlife conflict research that has shown that learning associated with negative reinforcement can be an effective way to reduce animal behaviors that lead to human-wildlife conflict (Much et al., 2018) and that guarding is perceived as an effective non-lethal countermeasure (for carnivores; Scasta et al., 2017; Young & Sarmento, 2024). Floodlights and speakers that played predator calls both reduced farm invasion, but only when used as intended. Taken together, these data suggest that giraffe-farmer conflict can be reduced by a variety of pathways, but most effectively when best practices are consistently used in conjunction.

Local weather patterns influence crop raiding by giraffes—surveys of both giraffe populations and farmers indicate that crop raiding by giraffes primarily occurs in dry seasons, which coincides with increased abundances of giraffes near riverfront farmlands (supporting our Hypothesis 1; Fig. 3). Although most wildlife are naturally afraid of humans and therefore avoid spaces with high levels of human activity (Carter et al., 2012), there are situations when wildlife cannot avoid being in human-occupied areas even at peak levels of activity. Scarcity of resources, for example, may sometimes outweigh the need for safety (Oates et al., 2019; Calhoun et al., 2025). Weather patterns influence availability of vegetative resources within landscapes and although the trees that giraffes primarily feed on are deeply rooted and slow at being affected by short term weather changes, drought has become increasingly common both in East Africa and globally (Ayugi et al., 2022; Calhoun et al., 2025; Gebrechorkos et al., 2025). Drought not only reduces available forage through leaf shedding (most local tree species are drought-deciduous), but also through alteration of leaves’ chemical composition in drought-resistant plants (Furstenburg & van Hoven, 1994; Caister et al., 2003). Riverine vegetation sustained by locally elevated water tables has therefore become important for the conservation of giraffes in arid environments (Caister et al., 2003). In our study area, riverine vegetation includes food crops (especially mangoes) that are increasingly becoming an often-used food source for giraffes during the dry season. Seasonally deploying mitigation measures during dry seasons may increase the logistical feasibility and cost-effectiveness of employing mitigation measures to prevent crop depredation.

Mangoes are significantly associated with farm invasion by giraffes (Hypothesis 2). Because different herbivore species eat different parts of plants at different development stages, crop type often influences the extent of crop raiding, the wildlife species involved in it, and when it occurs. Non-palatable crops have therefore been proposed as a way of reducing crop raiding (Seiler & Robbins, 2016; Matsika et al., 2023; Yogendra et al., 2024). Giraffes are selective feeders that select young shoots, pods, seeds, and flowers when available (Furstenburg & van Hoven, 1994; Fleming et al., 2006), which may increase conflict because giraffes overlap with farmland during dry seasons when mangoes flower (Cavalcante, 2022). Countermeasures targeted at preventing giraffe invasions will be most effective if they target mango plantations, and replacing mango trees with citrus (e.g., lemon and lime) trees may reduce farmer-giraffe conflict in high-priority areas for giraffe conservation such as Garissa. However, future studies should examine the roles of multiple crops grown together and if certain crops are reliable deterrents, as we did not explicitly test these factors. In addition, most of the farms (11 out of 12) that did not experience invasion and did not grow mangoes were spatially clustered near Sankuri (11 farms; 29% of farms surveyed in Sankuri). This suggests that there may be collective benefits when farms in an area are robust to invasion, as giraffes may collectively learn that the area does not provide food resources. In addition, interacting climatic and economic factors that lead to crop decisions in an area are likely to influence this pattern. Although we did not explicitly test this, this is a potential dimension of countermeasure adoption that should be explored.

Countermeasures that targeted giraffe anatomy by creating physical barriers were perceived as ineffective, but countermeasures targeted at behavior showed potential when properly and consistently used (Hypothesis 3A). Large mammals like giraffes can break or jump over physical barriers like fences. For fences to successfully deter giraffes, they must either be tall enough for giraffes to not simply walk over them, stout enough that giraffes cannot break through them, or electrified. In our study area, most of the fences were made from piles of small thorny branches of mesquite, *Prosopis juliflora,* cleared during farm preparation (Fig. S1). Where fences were reported to be effective, they were made of wire, concrete blocks, or very large, fresh piles of branches. Deployment of better fence technology would likely increase the efficacy of fences but would also come with high costs that may not be feasible for most farmers in the area.

### Regardless, the farmers appeared to favor fencing as a countermeasure, particularly improvement to wire fences

Human guards were the most effective countermeasure. In most cases, human guards consisted of full-time farm attendants and were only available for those who could afford them or if the owner lived at the farm and served as the full-time guard. Because it was expected that guards work on the farm the next morning, human guards were intermittently active and relied on other methods such as guard dogs to alert them of invasion by giraffes. Although human guards were reported as the most effective countermeasure used by farmers to reduce farm invasion by giraffe (Fig. S3), they were not the leading countermeasure farmers were willing to try because human guards are expensive and require consistent effort to maintain. Farmers were instead most interested in passive countermeasures such as fences, fire, sound devices, flashlights and guard dogs. In addition to guard dogs—which were mainly used to alert human guards—fire, sound devices, and flashlights were used to simulate presence of humans on the farm. When fire was used as a countermeasure, it was lit at known entry points for giraffes. Fire was effective as long as it kept burning, but this also posed a risk of spreading onto the piles of branches used as fences. Giraffes were in the long run reported to become tolerant of fire. Sound devices and flashlights were closely associated with human guards who use them while driving giraffes from the farms. Like fire, when used in isolation, giraffes became tolerant of these methods. It should be noted that for religious reasons, few farmers used guard dogs, and guard dogs were often associated with farm attendants that were not Muslims.

In our study area, reports from farmers aligned with giraffe survey data that indicated that farm invasion occurs primarily during the dry season (Hypothesis 4A). Contrary to our expectation, while farmers perceived that reducing farm invasion using physical barriers was improbable, they also largely supported the idea of using improved fences (e.g., wire). This may be explained by the cost of the most effective countermeasure (human guard), but contrary to our expectations, the cost of a countermeasure was rarely reported as an important factor influencing willingness to try a new countermeasure (Hypothesis 4B; Fig. 7). Local availability, ease of use, and seeing neighbors using a countermeasure successfully were the leading factors that determined willingness to try a new countermeasure. Most farmers in our study area were also members of farm associations and shared knowledge on what to plant and use as countermeasures. While farmer perceptions were largely aligned with observed giraffe behavior, decisions on what countermeasures to implement could be improved, though they will largely depend on local farmers’ willingness to implement best practices and share information with neighbors.

Because countermeasures that targeted giraffe behavior (e.g., causing or exploiting fear of humans) were more effective than physical barrier-based countermeasures (e.g., makeshift fences), we tested the efficacy of affordable floodlights and sound devices as additional possible countermeasures. Although they were not effective 100% of the time, when used correctly and consistently, both light and sound devices meaningfully reduced farm invasions by giraffe (Hypothesis 3B). Farmers were initially skeptical about the effectiveness of the sound and light devices we provided. Initial success led to more farmers volunteering their farms for these countermeasures to be tested. Because we could not provide every farmer these countermeasures, farmers registered to farm associations shared the lights with other farmers. With limited motion sensor range, giraffes would invade the farm without triggering the lights. Some farmers reported lights and speakers being stolen, while other farmers used lights in their houses and speakers to play music rather than deter giraffes. While floodlights showed the most potential for reducing farm invasion, a single solar floodlight costs ca. $10, and the cost for installation increased with the size of the farm due to the limited range of their motion sensors (ca. 5m) requiring that an entire farm be surrounded with floodlights at ca. 9m intervals. Light was therefore more suitable for small farms. Sound devices, on the other hand, were more suitable for large farms. Farmers expressed concerns that lion roars were unsettling for livestock and were sometimes stolen, leading to some farmers to use them inconsistently. Further experiments could help corroborate the efficacy of these countermeasures, determine the extent to which giraffes eventually habituate to these countermeasures, and develop methods to increase adherence to best practices.

Crop raiding by giraffes and resulting retaliatory measures from farmers (e.g., snaring, spears and machete wounds) are an emerging issue in giraffe conservation as more lands are converted to farmland to meet the increased demand for food of a growing human population living in a changing climate (Abrahms et al., 2023; Fagan et al., 2022). This has also been facilitated by changes in the land-tenure system in Kenya, which has shifted from predominately communal ownership to more private individual ownership (Mwangi, 2024; Di Matteo, 2025; Ali et al., 2026). This has made nomadic pastoralism harder to sustain and the previously nomadic community in our study area is becoming more sedentary and practicing irrigated farming along the only permanent source of water, the Tana River. Similar trends are occurring in other parts of the country (Di Matteo, 2025) and elsewhere in Africa. Effective giraffe conservation will, therefore, depend on increasing our understanding of the multiple drivers of this emerging source of human-wildlife conflict, and developing interventions that are supported by both communities and governments (Torres et al., 2018).

## Acknowledgments

We would like to thank the Conservation Leadership Programme for their generous financial support to deliver this project (award #01423421). Many thanks to all the Bour-algy Giraffe Sanctuary rangers, especially Siyat Haji for accompanying, and helping throughout the project via his knowledge of local farmers. A special thanks to the Somali Giraffe Project and the team for hosting this project while conducting our fieldwork. An honorable mention to the Face of African Conservation Ecology group, a peer-to-peer mentorship for support while brainstorming about feasibility of the project in pursuit of grant and during field implementation. Additional funding for this research was provided by the American Society of Mammalogists through an African Research Fellowship, Society for Conservation Biology, and the School of Natural Resources and the Environment at the University of Arizona.

